# Chromatin organization changes during the establishment and maintenance of the postmitotic state

**DOI:** 10.1101/184994

**Authors:** Yiqin Ma, Laura Buttitta

## Abstract

**Background:** Genome organization changes during development as cells differentiate. Chromatin motion becomes increasingly constrained and heterochromatin clusters as cells become restricted in their developmental potential. These changes coincide with slowing of the cell cycle, which can also influence chromatin organization and dynamics. Terminal differentiation is often coupled with permanent exit from the cell cycle and existing data suggests a close relationship between a repressive chromatin structure and silencing of the cell cycle in postmitotic cells. Here we examine the relationship between chromatin organization, terminal differentiation and cell cycle exit.

**Results:** We focused our studies on the *Drosophila* wing, where epithelial cells transition from active proliferation to a postmitotic state in a temporally controlled manner. We find there are two stages of G_0_ in this tissue, a flexible G_0_ period where cells can be induced to re-enter the cell cycle under specific genetic manipulations and a state we call “robust”, where cells become strongly refractory to cell cycle re-entry. Compromising the flexible G_0_ by driving ectopic expression of cell cycle activators causes a global disruption of the clustering of heterochromatin-associated histone modifications such as H3K27 trimethylation and H3K9 trimethylation, as well as their associated repressors, Polycomb and heterochromatin protein 1(HP1). However, this disruption is reversible. When cells enter a robust G_0_ state, even in the presence of ectopic cell cycle activity, clustering of heterochromatin associated modifications are restored. If cell cycle exit is bypassed, cells in the wing continue to terminally differentiate, but heterochromatin clustering is severely disrupted. Heterochromatin-dependent gene silencing does not appear to be required for cell cycle exit, as compromising the H3K27 methyltransferase *Enhancer of zeste*, and/or HP1 cannot prevent the robust cell cycle exit, even in the face of normally oncogenic cell cycle activities.

**Conclusions:** Heterochromatin clustering during terminal differentiation is a consequence of cell cycle exit, rather than differentiation. Compromising heterochromatin-dependent gene silencing does not disrupt cell cycle exit.

## Background

Cellular differentiation is the acquisition of cell-type specific characteristics, driven by changes in gene expression. Changes in gene expression are largely controlled by transcription factors, which can be facilitated or impeded by chromatin modifications, binding site accessibility and chromatin organization. A reciprocal relationship exists between chromatin organization, modification and gene expression, and several studies have shown that chromatin organization and modifications can change during differentiation. For example, during neural differentiation silenced genes move to repressive compartments in the nucleus [1–3]. In certain contexts of differentiation global nuclear compartments can become dramatically re-organized to facilitate specialized functions [4]. At a more local level, chromatin modifiers can be recruited to specific genes involved in differentiation to facilitate their expression and limit the expression of genes involved in other cell-type programs that must be kept off [5]. Thus dynamic changes in chromatin organization and modification can have critical consequences on proper differentiation during development.

There is also an intimate relationship between the cell cycle and chromatin organization and modifications. Chromatin in actively cycling cells is highly dynamic. During S-phase, new histones are incorporated onto nascent DNA requiring re-establishment of histone modifications [6]. During mitosis, nuclear organization including intra and inter-chromosomal contacts are lost and many chromatin modifiers are ejected from chromatin to facilitate proper chromosome condensation and segregation [7, 8]. In addition the activity of histone modifiers can be regulated in a cell cycle-dependent manner [9–14]. During differentiation cells often transition from rapid proliferation to slower cycling, which can be followed by cell cycle exit or entry into G_0_ coordinated with terminal differentiation. Thus the modification and organization of chromatin in the nucleus can be impacted by the differentiation process itself, but also by the changes in cell cycle dynamics during differentiation. For example, chromatin compacts and heterochromatin clusters as cells in the embryo cycle more slowly and become lineage restricted [15]. In *Drosophila* loci within constitutive heterochromatin show increased association in terminally differentiated postmitotic cells [16] and facultative heterochromatin-forming Polycomb bodies cluster as cells differentiate and the cell cycle slows during embryogenesis [17]. Methods such as inducing developmental arrest have been used in attempt to disentangle the influence of cell cycle changes from differentiation process [16], but these approaches cannot fully uncouple terminal differentiation from the accompanying cell cycle exit and it has remained unclear whether changes in heterochromatin clustering and dynamics are due to differentiation, the accompanying cell cycle changes, or both. The influence of cell cycle changes during differentiation adds a layer of complexity to our understanding of the relationship between chromatin organization and modifications and differentiation.

Here we directly address the relationship between heterochromatin organization, chromatin modification and cell cycle exit using the temporally controlled cell cycle exit in the *Drosophila* wing [18–20]. In our experiments, we take advantage of tools that can effectively uncouple cell cycle exit and differentiation to ask whether heterochromatin clustering is a consequence of cell cycle exit or differentiation. In addition we examine changes in chromatin modifications caused by the delay of cell cycle exit and examine the impact of disrupting heterochromatin-dependent gene silencing on cell cycle exit.

## Methods

### Fly stocks and genetics

#### Disruption of G0 in the posterior wing

*w*/*y*, *w*, *hs*-*FLP*; *en*-*GAL4*, *UAS*-*GFP*/ *UAS*-*E2F1*, *UAS*-*DP*; *tub*-*gal80TS*/ +

*w*/*y*, *w*, *hs*-*FLP*; *en*-*GAL4*, *UAS*-*GFP*/ *UAS*-*CycD*, *UAS*-*Cdk4*; *tub*-*gal80TS*/ *UAS*-*E2F1*, *UAS*-*DP*

*w*/*y*, *w*, *hs*-*FLP*; *en*-*GAL4*, *UAS*-*GFP*/ +; *tub*-*gal80TS*/ *UAS*-*CycE*, *UAS*-*Cdk2*

*w*/*y*, *w*, *hs*-*FLP*; *en*-*GAL4*, *UAS*-*GFP*/ *UAS*-*CycD*, *UAS*-*Cdk4*; *tub*-*gal80TS*/ +

*w*/*y*, *w*, *hs*-*FLP*; *en*-*GAL4*, *UAS*-*GFP*/ *UAS*-*CycA*; *tub*-*gal80TS*/ +

#### Disruption of G0 in clones

*w*/*y*, *w*, *hs*-*FLP*; *tub*>*CD2*>*GAL4*, *UAS*-*GFP*/ *UAS*-*CycD*, *UAS*-*Cdk4*; *tub*-*gal80TS*/ *UAS*-*E2F1*, *UAS*-*DP*

#### Disruption of H3K27me3

*w*/*y*, *v*; *en*-*GAL4*, *UAS*-*GFP*/ +; *tub*-*gal80TS*/ *UAS*-*E*(*z*)^*RNAi*^ (*Bloomington 33659*)

#### Disruption of HP1

*w*/*y*, *v*; *en*-*GAL4*, *UAS*-*GFP*/ +; *tub*-*gal80TS*/ *UAS*-*Su*(*var*)*205*^*RNAi*^ (*Bloomington 33400*)

#### Disruption of HP1 with Y10C

*w*/*w*, *Y10C*; *en*-*Gal4*, *UAS*-*RFP*/ +; +/ *UAS*-*Su*(*var*)*205*^*RNAi*^ (*Bloomington 33400*)

All the crosses containing gal80^TS^ were maintained in 18°C to suppress Gal4 in early development. To disrupt G_0_ with cell cycle regulators, white prepupae were collected and shifted to 28°C to indicated time points. For *E(z)* knockdown experiments, L3 larva were shifted from 18°C to 28°C to induce *E(z)* RNAi. For *HP1* knockdown, crosses were kept in 28°C after egg laying (AEL). For clonal expression of cell cycle regulators, animals were heat shock in 37°C for 8 minutes during 48-72h AEL, and then kept in 18°C. White prepupae were collected and shifted to 28°C to indicated time points. All timings are adjusted according to the equivalent development at 25°C as described previously [18].

### Immunostaining

Imaginal discs or pupal wings were dissected in 1x PBS, and fixed in 4% paraformaldehyde /1x PBS for 30 minutes. Samples were washed twice in 1x PBS, 0.1% Triton X, 10 min each, and incubated in PAT (1XPBS, 0.1% Triton X-100 and 1% BSA) for 10 mins for larval tissues and 3 x 20 mins for pupal tissues. Samples were then incubated with primary antibodies for 4 hours or 4°C overnight followed by 3 washes and secondary antibodies at room temperature for 4 hours or 4°C overnight. Primary antibodies used in this study include: Anti-phospho-Ser10 histone H3, 1:2000 rabbit (Millipore #06-570) or mouse (Cell Signaling #9706); Anti-GFP, 1:1000 chicken (Life Technologies A10262) or 1:1000 rabbit (Life Technologies A11122); Anti-pH2Av, 1:100 mouse (DSHB, UNC93-5.2.1); Anti-H3K27me3, 1:500 rabbit (Millipore #07-449); Anti-HP1, 1:250 mouse (DSHB, C1A9); Anti-H2Av, 1:500 rabbit (Active Motif #39715); Anti-H3, 1:500 mouse (Cell Signaling #3638); Anti-H3ac, 1:500 rabbit (Millipore #06-599); Anti-H3K4me3, 1:500 rabbit (Millipore #07-473); Anti-H3K9me3, 1:500 rabbit (Millipore #07-523) or (Active Motif #39161); Anti-H3K27ac, 1:500 rabbit (Abcam ab4729); Anti-H4ac, 1:500 rabbit (Millipore #06-866); Anti-H4K16ac, 1:500 rabbit (Millipore #07-329); Anti-H4K20me3, 1:500 mouse (Abcam ab78517); Anti-E2F, 1:500 guinea pig (kindly provided by Dr. Terry L. Orr-Weaver); Anti-Ubx, 1:250 mouse (DSHB, FP3.38); Anti-D1, 1:200 guinea pig (kindly provided by Dr. Yukiko Yamashita). DNA was labeled by 1 ug/ml DAPI in 1× PBS, 0.1% Triton X for 10 min. F-actin was stained using 1:100 rhodamine–phalloidin (Invitrogen; R415) in 1x PBS for 4 hours.

### Microscopy and Image quantification

Images were taken with a 100x oil objective on a Leica SP5 confocal with a system optimized z-section of 0.13 μm. 3-D reconstructions were performed using the “3D viewer” function in Leica LAS AF software. Images of whole pupal wings in Fig. 2 and 7 were obtained using a Leica DMI6000B epifluorescence system. All adjustments of brightness or contrast were applied to the entire image in Adobe Photoshop and performed equally with equal threshold values across control and experiment samples.

For integrated intensity quantifications, we used maximum projections of 12 continuous z-sections of confocal images. We developed a toolkit in Matlab (Release 2015b) that automatically segments nuclei and foci within nuclei and integrates the pixel intensities with the help of the Advocacy and Research Support, U.Michigan LSA-IT. To identify nuclei. images were smoothed using a circular averaging filter through the fspecial and imfilter function of Matlab. Next a watershed algorithm was applied to segment nuclei from the background and nuclei were masked using local maxima with an h-maxima transform. Thresholds were manually set and checked for each image to accurately delineate nuclei. GFP positive vs. negative was established using an intensity threshold for the GFP channel. Integrated intensities for all nuclei were exported to Excel. Segmentation and measurement of foci followed a similar process for foci within the defined nuclear regions. In brief, foci were segmented using a watershed algorithm, then further measured for pixel intensity and number, which was used for foci area and intensity measurements.

**Fig. 2.**
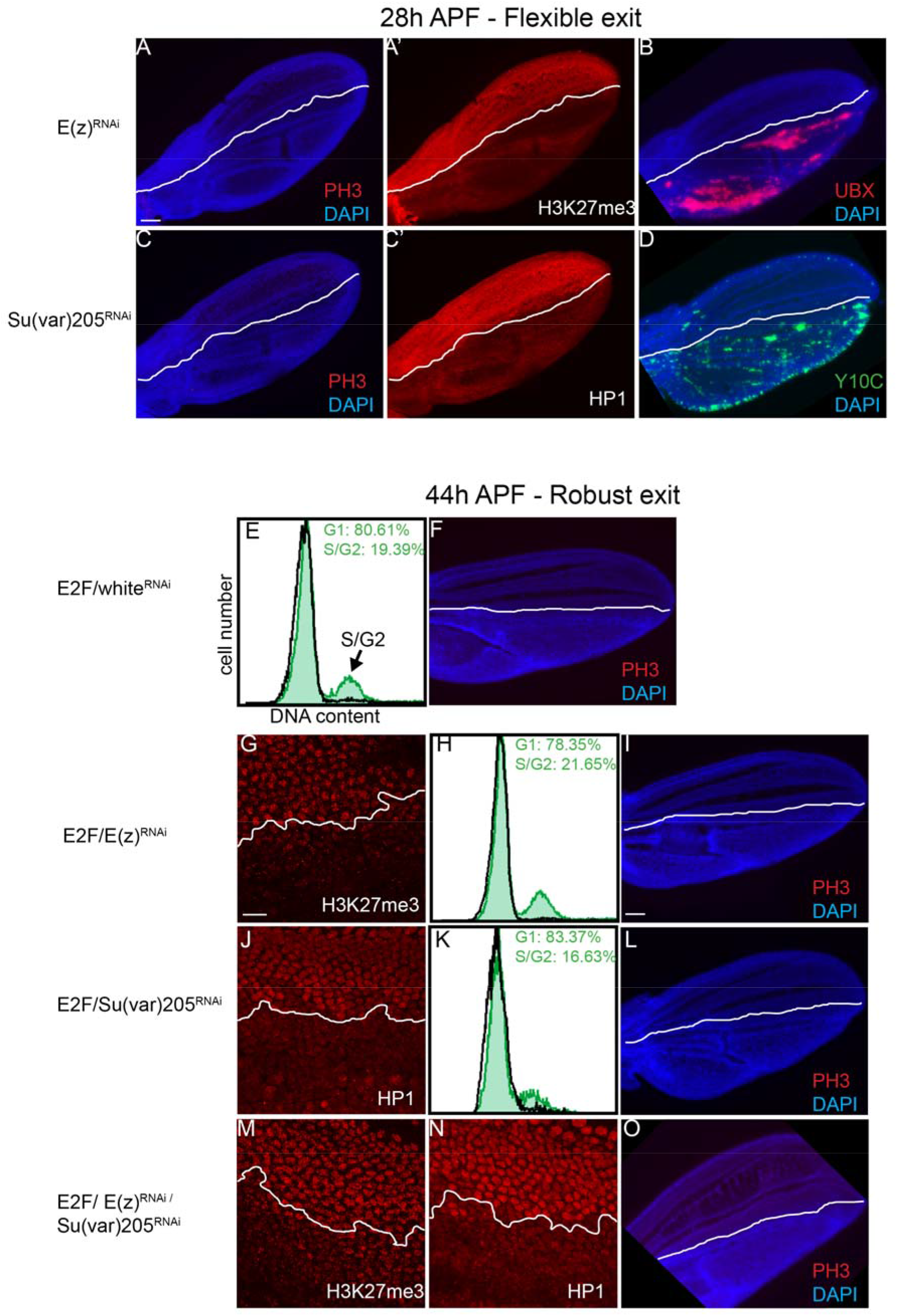
Heterochromatin-dependent gene silencing is not required for cell cycle exit. (A-B) RNAi to *E(z)* was expressed in the posterior wing from the L3 stage until the indicated timepoints in metamorphosis. Postmitotic wings at 26-28h were examined for mitoses as indicated by phospho-Ser10-histone H3, PH3 (A), H3K27me3 (A’) and de-repression of Ubx (B). (C) RNAi to *Su(var)205* (the gene encoding HP1) was overexpressed in the posterior wing and postmitotic tissues were immunostained for PH3 and HP1 (C’). These conditions led to loss of HP1 and disrupted heterochromatin-mediated silencing of the Y10C reporter (D). Control RNAi (to the *white* gene), *E(z)* and/or *Su(var)205* was expressed in the posterior wing in combination with E2F from the start of metamorphosis. Postmitotic wings at 42-44h were dissected and examined for H3K27me3 (G, M), HP1 (J, N) and PH3 (F, I, L, O). (E, H, K) Flow cytometry was also performed to measure cells that enter S and G2 phases. Green trace indicates cells from the posterior wing expressing the indicated transgenes. Black trace: control non-expressing anterior wing cells. Reduced heterochromatin gene silencing does not compromise G0 even in the presence of high E2F activity. Scale bars=50 μm A-D,F,I,L and O; 10 μm G,J,M and N.

**Fig. 7.**
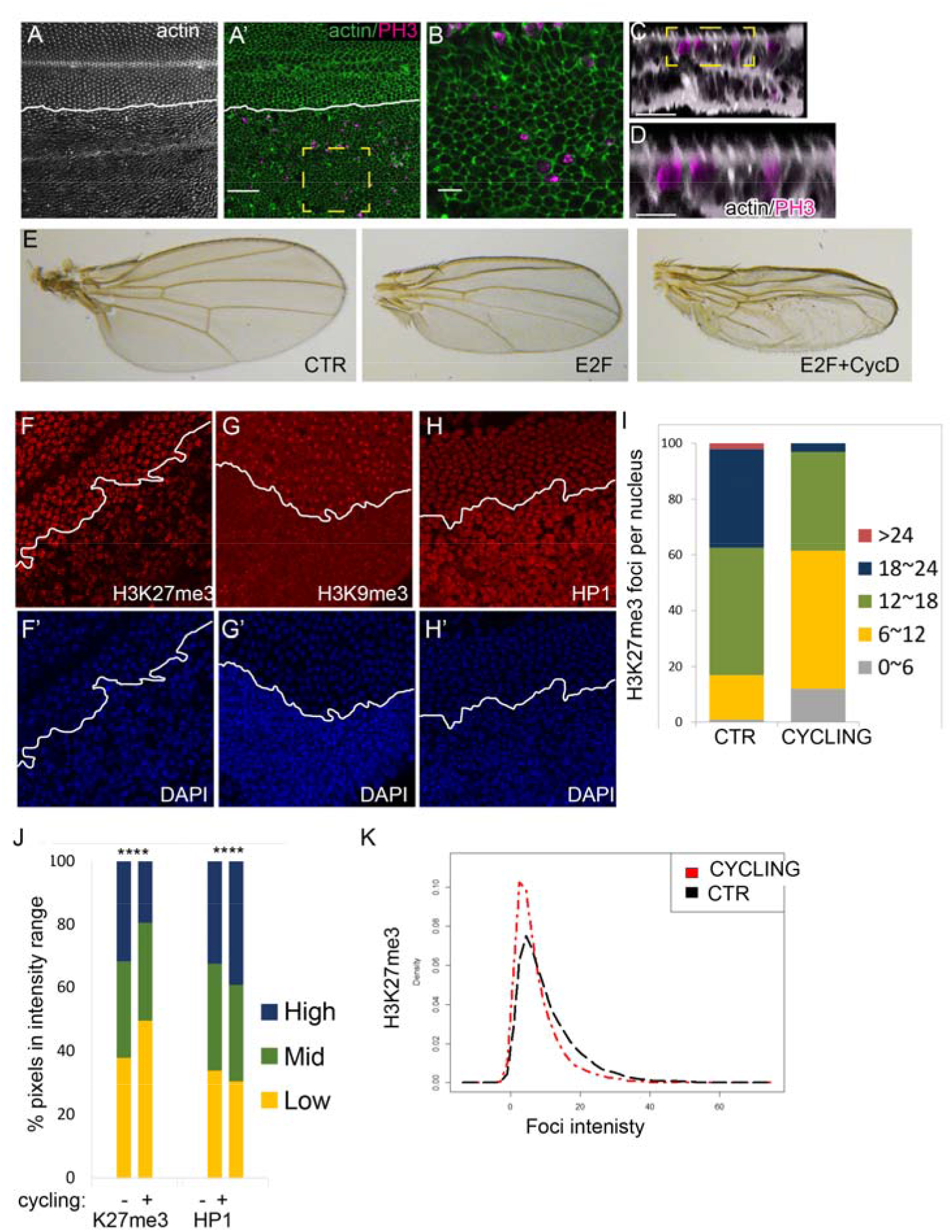
Heterochromatin clustering during terminal differentiation is a consequence of cell cycle exit. CycD, Cdk4 and E2F were co-expressed in the posterior wing to bypass robust cell cycle exit without preventing terminal differentiation. The anterior-posterior boundary is indicated by a white line. (A-D) Pupal wings at 42-44h were dissected stained for actin and PH3 to label mitoses. Mitoses are evident in cells generating wings hairs, a hallmark of wing terminal differentiation (E) and the wings generate intact adult wing cuticle. C and D show optical cross sections (x/z) of wings to reveal PH3 and actin-rich hairs in the same section. (F-K) CycD, Cdk4 and E2F expression in the posterior wing prevents G0 entry and disrupts proper localization of heterochromatin associated histone modifications and HP1. (J) The distribution of staining intensity in 474-1191 nuclei, binned into three ranges is shown. The reduced foci number (I) and intensity (K) indicates compromised clustering of H3K27me3 containing chromatin when entry into G0 is prevented. P-values were determined by an unpaired t-test; ****<0.0001. Scale bars=10 μm in A, 5 μm in B and 2.5 μm in D.

### Fluorescent *in situ* Hybridization (FISH)

Alexa-488 probes against the rDNA Internal Transcribed Spacer (ITS) region and Cy3 probes against AACAC repetitive satellite sequences were kindly provided by Dr. Yukiko Yamashita. FISH was performed on larval and pupal wings as described [21]. In brief, fixed tissues were treated with 2 mg/ml Rnase A in 1x PBS, 0.1% Triton X at 37°C for 10 min, and rinsed in 2x SSC/1mM EDTA/0.2%Tween 20. Then tissues were incubated in 2x SSC/1mM EDTA/0.2%Tween 20 solution with increasing formamide concentration from 20%, 40% to 50% for 15 min to 30 min. Finally, tissues were incubated in 100 μl hybridization solution with 50 μl formamide, 20 μl 50% dextran sulfate, 20 μl 2X SSC/1mM EDTA/0.2%Tween 20 and 10 μl of probe at μg/mL for 15 min at 91°C, and left at 37°C overnight. Quantification of size for rDNA loci area was carried out using our customized Matlab toolkit. For the quantification of AACAC satellite to the chromocenter, we used a single 0.13 μm z-section with the strongest FISH signal and measured the relative distance of the center of the FISH signal to the brightest Dapi-stained region and corrected for the total nuclear radius using the Leica LAS AF software.

### Flow cytometry

FACS was performed on dissociated wings to measure DNA content on an Attune Cytometer (Life Technologies) as described [22].

### RNA interference

Kc167 cells were kindly provided by Dr. K. Cadigan and cultured as described [23]. For RNA interference, cells were placed with concentration of 1 million/ml and starved in serum free medium with 10 ug/ml double-strand RNA (dsRNA) for 4-6 hours, then 10% serum medium was added to the culture and cells were collected for staining 3 days after serum medium addition. dsRNA was synthesized with T7 Megascript Kit (Ambion). T7 primers used in this study:

T7-Wee-fwd, TAATACGACTCACTATAGGGATGACTTTGACAAGGACAC; T7-Wee-rev, TAATACGACTCACTATAGGATCTAGTCGATTGACGCATT; T7-Myt1-fwd, TAATACGACTCACTATAGGAATTGCACGACGACAAACAC; T7-Myt1-rev, TAATACGACTCACTATAGGTGTCCAGATGGATGAGATTC; T7-Myt1-fwd2, TAATACGACTCACTATAGGACAACAATCTGAACCGAAGC; T7-Myt1-rev2, TAATACGACTCACTATAGGTGGAGCCATATACCTCGAAT;

### Western Blots

Western Blots were performed on staged fly wings using BioRad TGX precast 4-20% gels and high sensitivity ECL reagents (Thermo) to detect HRP conjugated secondary antibodies [24]. Mouse anti-α-tubulin (1:1000, DSHB, AA4.3) was used as a loading control. Blot signals were detected and quantified with FluorChem M digital system from Protein Simple.

## Results

### Heterochromatin clusters as proliferation slows and cells differentiate

The impact of the cell cycle on heterochromatin clustering during cellular differentiation has not been resolved. Specifically, how the transition from a proliferative to a postmitotic state impacts global chromatin organization in *Drosophila* is unclear. To examine this, we immunostained for various chromatin marks and chromatin binding proteins in wild-type *Drosophila* wings at three stages with distinct proliferation parameters. We examined quickly proliferating second instar larval (L2) wings, slowly proliferative wandering third instar larva (L3) and post-mitotic 28h pupal wings (Fig.1A). Cells of the L2 wing region examined have a cell doubling time (CDT) of about 10h, while cells of the same region in L3 wings have a longer CDT of 15h. By 28h after puparium formation (APF) during metamorphosis, cells of the wing blade have entered G0 and are permanently postmitotic [18, 25, 26]. We examined the histone modification H3K27me3 associated with facultative heterochromatin, H3K9me3, HP1 and the AT rich repetitive sequence binding protein D1 associated with constitutive heterochromatin and the euchromatin-associated modification H3K4me3 (Fig. 1A). The immunofluorescence (IF) signals for H3K27me3, H3K9me3 and D1 were weakest at the L2 stage, but increased at the L3 and pupal stages and clustered into larger and more intense, distinct foci in the slower cycling tissues (Fig. 1A). In *Drosophila* cells, the chromocenter, containing constitutive heterochromatin such as clustered centromeres, can be easily visualized as a DAPI-bright region within the nucleus [27]. We confirmed the co-localization of the chromocenter with D1 staining, which binds centromeric satellite repeats, and also co-localized with the centromeric histone Cenp-A (not shown) [28]. H3K9Me3 and HP1 label heterochromatin foci partially overlapping and adjacent to the DAPI-bright region [29]. H3K27Me3 labels distinct foci throughout the nucleus associated with facultative heterochromatin, and represents Polycomb repressive complex 2 (PRC2) binding and formation of Polycomb group (PcG) clusters or foci [30–32]. By contrast, H3K4Me3 broadly localizes throughout the chromatin, does not form distinct foci, and is excluded from the centromeric and pericentromeric regions (Fig1A).

**Fig.1.**
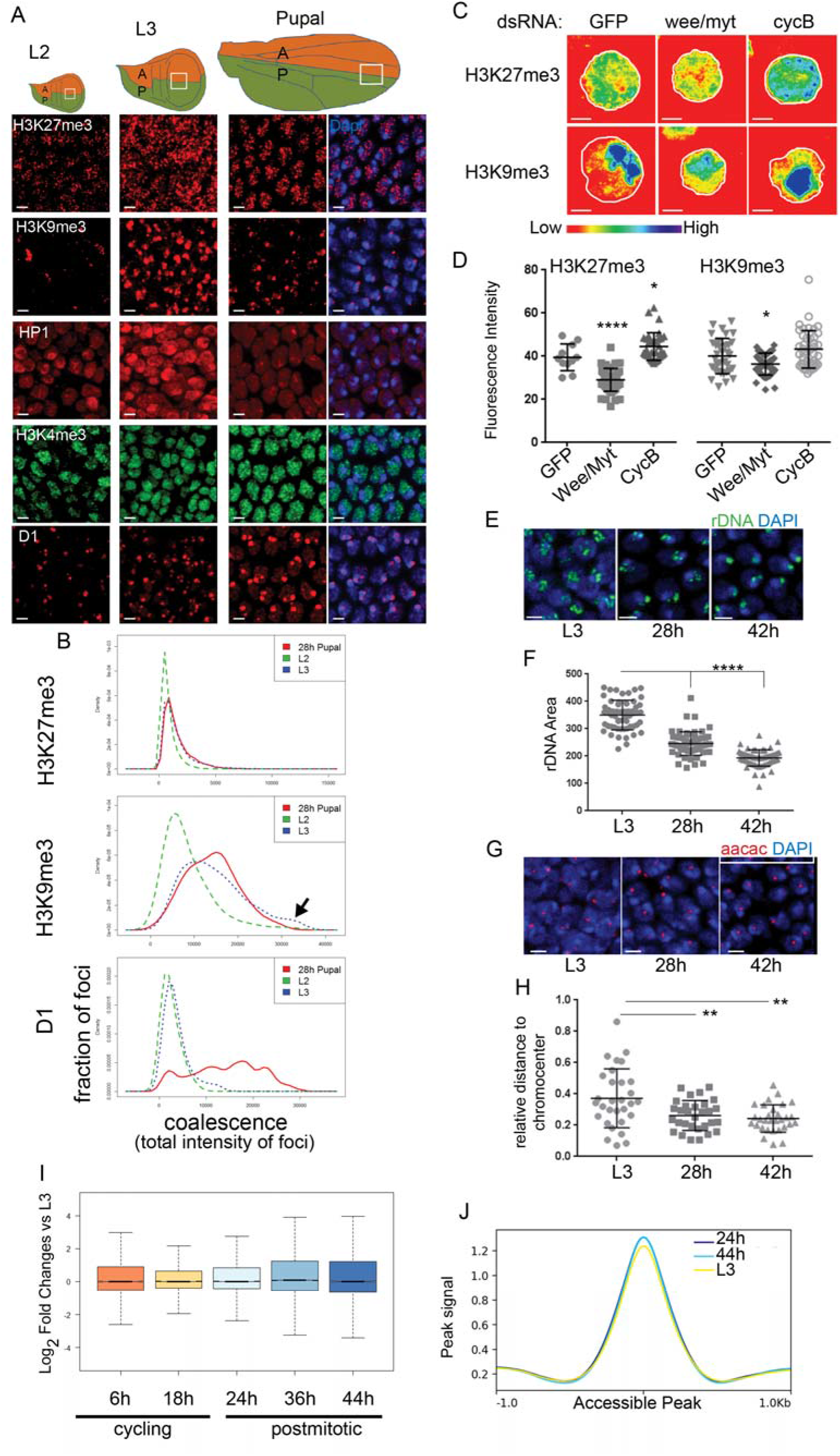
Heterochromatin clustering increases as the cell cycle slows and cells differentiate. (A) Wings of the indicated developmental stages were immunostained for the indicated chromatin modifications and chromatin binding proteins. (B) As the cell cycle slows down and cells differentiate, the distribution of heterochromatin-associated foci shift toward larger, brighter foci indicating increased clustering. Coalescence is quantified as the total intensity of individual focus within 129-448 nuclei at each developmental stage. (C,D) KC cells treated with dsRNA against GFP, *wee/myt* and *cycB* were immunostained for the indicated chromatin modifications and fluorescence intensity was quantified. (E,F) Fluorescent in situ hybridization (FISH) against the rDNA ITS region was performed on wings of the indicated stages. rDNA foci coalesce and condense in postmitotic cells. (G, H) FISH against the AACAC pericentromeric satellite repeats was performed on wings of the indicated stages and the distance to the center of the DAPI-bright chromocenter was measured. The distance decreases in postmitotic cells indicating increased condensation of heterochromatin. (I) A box plot of the RNA log2-fold changes compared to proliferative L3 for each timepoint is shown. (J) A line plot of average FAIRE-seq signal across all accessible chromatin for the indicated stages is shown. The accessibility of regulatory elements are similar in cycling and postmitotic wings. Scale bars=2 μm. P-values were determined by an unpaired t-test *<0.05, **<0.01, ***<0.001, ****<0.0001.

To automatically detect and measure heterochromatin foci parameters such as intensity and number for a large number of nuclei, we developed a custom MatLab App (described in supplemental methods) that uses DAPI staining to mask individual nuclei followed by foci segmentation and measurement. We measured clustering of heterochromatin foci as a function of the integrated intensity for each focus (the sum of intensities for all pixels in a focus) [17, 32]. This automated approach allowed us to examine the distribution of heterochromatin foci at a single cell level, across hundreds to thousands of nuclei, sampled from multiple wings for each experiment in an unbiased manner. We found that heterochromatin clustering increased as the cell cycle slowed and stopped during L3 and pupal stages (Fig1B). We noted a dramatic increase in H3K9Me3 and HP1 staining at the L3 stage, which may reflect a developmentally controlled stage-specific increase in this modification/reader pair.

To distinguish whether an increase in heterochromatin clustering is due to changes in the cell cycle, we turned to *Drosophila* cell culture. In *Drosophila* Kc cells, the overall cell doubling time is controlled by the negative and positive regulators of the G2/M transition Wee/Myt and CycB respectively [23]. We sped up the cell cycle by reducing Wee/Myt1 activity via RNAi or slowed the cell cycle using RNAi to *cycB*. Slowing the cell cycle increased the clustering and intensity of H3K27Me3 and H3K9Me3 compared to controls exposed to RNAi to GFP (Fig1C-D).

The increased clustering of heterochromatin could be due to chromatin condensation and compaction. To examine chromatin condensation we performed Fluorescent *In Situ* Hybridization (FISH) using probes against the internal transcribed spacer region between the 18S RNA and 28S rDNA loci, which are tandemly repeated on the X, and measured the total rDNA area before and after cell cycle exit in the wing [33, 34]. In proliferating L3 wings the rDNA is extended. The rDNA becomes more compact as cells enter G0 at 24-28h APF and condense further as G0 is maintained at 42h APF (Fig. 1E-F). The changes in the rDNA locus suggest chromatin condensation increases in prolonged G0. To verify that compaction is not specific to the rDNA locus on the X, we also performed FISH to the pericentromeric satellite repeat AACAC on chromosome II and measured the distance of the signal to the chromocenter (Fig. 1G-H). The distance of the pericentromeric heterochromatin to the chromocenter also decreased suggesting that heterochromatin condensation, coalescence and compaction occurs throughout the nucleus after cell cycle exit.

An increase in chromatin clustering could be correlated with a global reduction in gene expression when cells become postmitotic [35]. To test whether global gene expression is reduced in postmitotic wings we examined an RNAseq timecourse of gene expression from proliferating to postmitotic stages [23]. We found the global gene expression levels to be similar in proliferating and postmitotic tissues (Fig. 1I). Next we compared the global changes in chromatin accessibility between proliferating and postmitotic wings through Formaldehyde-assisted Identification of Regulatory Elements [FAIRE]-seq [36] (Fig. 1J). Consistent with the global gene expression profile, we found no obvious changes in the average level of chromatin accessibility in cycling vs. post mitotic tissue. This suggests that clustering of heterochromatin as cells exit the cell cycle is not due to global changes in gene expression during differentiation and cell cycle exit.

### Compromising heterochromatin-dependent gene silencing does not disrupt cell cycle exit

We have shown that heterochromatin clustering increases with entry into G0. Heterochromatin clustering is associated with increased target gene silencing [32], and has been suggested to repress cell cycle gene expression to facilitate cell cycle exit in mammalian muscle and neurons [37–39]. To test whether heterochromatin-dependent gene silencing promotes cell cycle exit in Drosophila wings, we compromised the H3K27me3 methyltransferase E(z) and/or the H3K9Me3 binding protein HP1. As E(z) and HP1 perform many functions during development, we turned to an inducible system with RNAi to alter gene function after embryogenesis. We used the *engrailed*-Gal4 driver with a temperature sensitive Gal80 (*en*-Gal4/Gal80^TS^) to turn on UAS-driven expression of dsRNAs to *E(z)* and *HP1* in the posterior wing from the early L1 and L3 stages respectively. We then dissected wings at 24-28h APF and stained for the mitotic marker phosphorylated phospho-Ser-10 histone H3 (PH3) to determine whether cells in the posterior wing delayed or bypassed cell cycle exit. We saw no effect of *E(z)* or *HP1* reduction on cell cycle exit despite a clear loss of H3K27Me3 and HP1 in the posterior wing (Fig. 2 A-C). We further confirmed that our knockdowns effectively compromised heterochromatin-dependent gene silencing in the wing, by examining de-repression of the Polycomb target Ultrabithorax (UBX) and the HP1-silenced Y10C GFP reporter (Fig. 2 B,D). Recent work has suggested Polycomb (Pc) can repress certain targets independent of *E(z)* [30]. We therefore also directly inhibited *Pc* by RNAi, but observed no effect on cell cycle exit despite de-repression of UBX in the wing (Supp. Fig. 2).

Compromising heterochromatin-dependent gene silencing does not disrupt or delay cell cycle exit on its own, but we wondered whether it may sensitize cells to other perturbations that compromise cell cycle exit. We have previously shown that activation of various cell cycle regulators including the cell cycle transcription factor complex

E2F/DP (hereafter referred to as E2F), can cause 1-2 extra cell cycles in the pupa wing between 24-36h APF followed by a delayed entry into G0 at 36 APF (Supp. Fig 3). We refer to the 24-36h APF period as flexible cell cycle exit or “flexible G0” which is followed by a more difficult to disrupt “robust G0” after 36h. We co-expressed E2F with RNAi to E(z) and/or HP1 to examine whether loss of heterochromatin-dependent gene silencing can further delay cell cycle exit in the presence of high E2F activity. However, inhibition of E(z), *HP1* or *E(z)+HP1* together did not further compromise cell cycle exit in the presence of high E2F activity (Fig. 2G-O). Altogether our results demonstrate that compromising heterochromatin-dependent gene silencing does not disrupt cell cycle exit in the *Drosophila* wing.

### Delaying cell cycle exit disrupts heterochromatin clustering and chromosome compaction

Constitutive and facultative heterochromatin clusters in post-mitotic wings. To examine whether compromising cell cycle exit affects clustering, we used the system described above to express E2F in the posterior pupal wing to drive 1-2 extra cell cycles and delay exit from 24 to 36h APF. We immunostained for the heterochromatin-associated histone modifications H3K27me3, H3K9me3 and H4K20me3 at 26-28h APF, a timepoint when E2F induces abundant mitoses in the posterior wing (Supp. Fig. 3). We compared the clustering of the chromatin marks in the unperturbed anterior to the posterior wing. When cell cycle exit is delayed, all three modifications appear more diffuse throughout the nucleus and heterochromatin clustering is disrupted (Fig. 3A-M). To determine whether E2F altered the total abundance of the modified histones, we performed semi-quantitative western blots on 28h pupal wings. With E2F expression, total levels of H3 were increased, consistent with additional S-phases leading to replication-coupled canonical histone production [40, 41]. However the ratio of modified H3 to total H3 was relatively unchanged or even slightly increased when cell cycle exit was delayed (Supp. Fig. 1). This may be because E2F activity also increases the expression of several PRC2 components (*E(z), esc, Su(z)12*) and *Su(var)3-9)* as well as several other histone modifying enzymes (Supp. Table 1), a feature conserved with mammalian E2Fs [42]. Thus, delaying cell cycle exit increases new histone production, but the histone modification rate is maintained by a coordinated increase in the expression of the modifying enzymes.

**Fig. 3.**
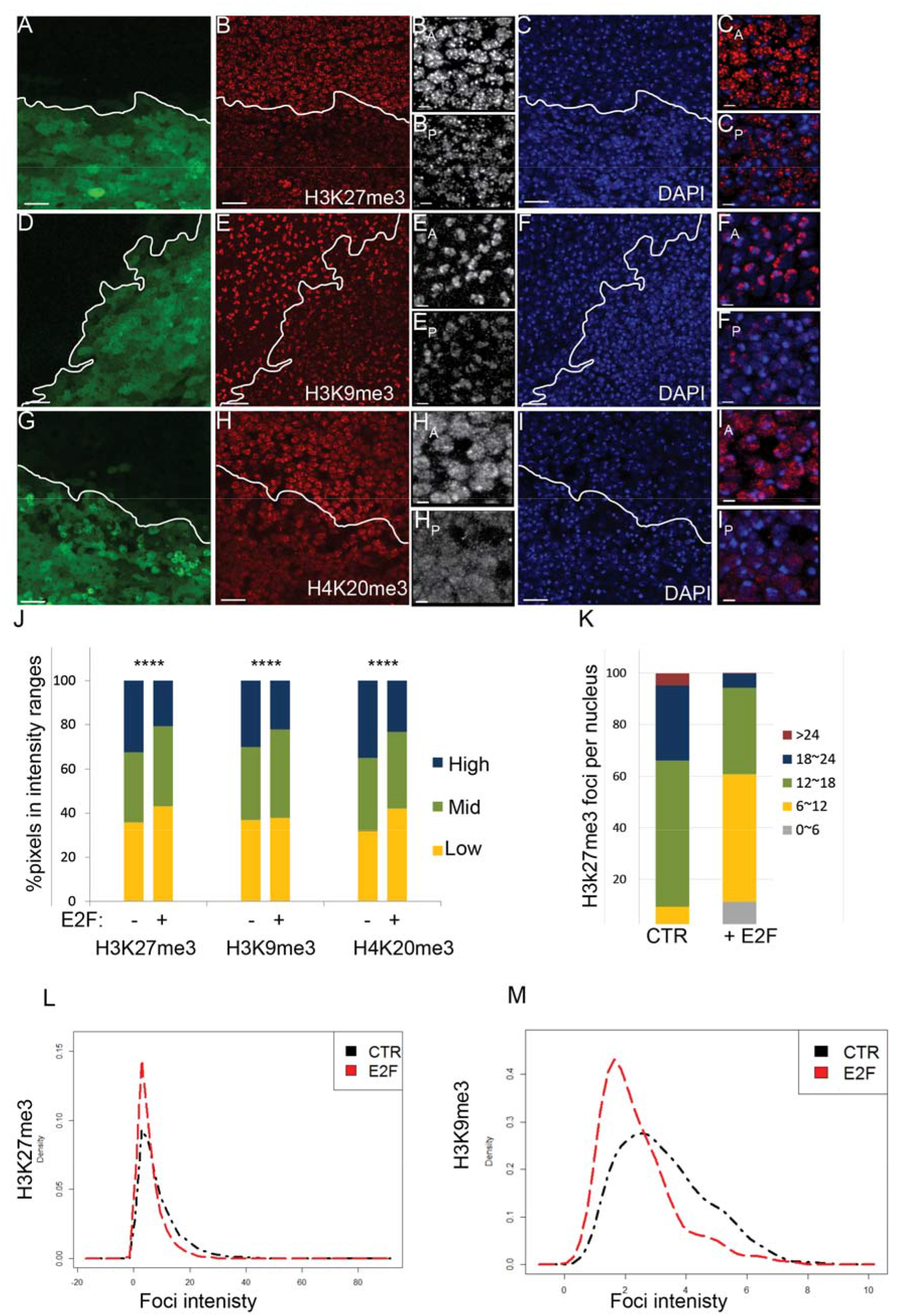
Heterochromatin clustering is disrupted when G0 is compromised. E2F was co expressed with GFP in the posterior wing (boundary indicated by a white line) from the start of metamorphosis (0h APF) to delay cell cycle exit. At 26-28h wings were dissected and immunostained for the indicated chromatin modifications and DAPI to label nuclei (A-I). (J-M) Fluorescence signal were measured for 485-848 nuclei for each chromatin modification. The distribution of overall fluorescence intensity (J), foci number per nucleus section (K) and individual foci intensity (L, M) all indicate that delaying cell cycle exit disrupts heterochromatin clustering in wing cell nuclei. Scale bars=10 μm in A-I except for anterior (A) and posterior (P) zoomed images where the bar = 2 μm (e.g. BA,BP). P-values were determined by an unpaired t-test. *<0.05, **<0.01, ***<0.001, ****<0.0001.

To determine whether delaying cell cycle exit also affected the localization of proteins associated with heterochromatin, we examined HP1, D1 and Polycomb using a Pc-GFP fusion protein [17]. We observed a more diffuse localization and a reduction in the clustering of these heterochromatin-associated proteins when cell cycle exit was compromised (Fig. 4A-M). This was also accompanied by a reduction in heterochromatin condensation, as assessed by the distance of the AACAC satellite to the chromocenter (Fig.4 N).

**Fig. 4.**
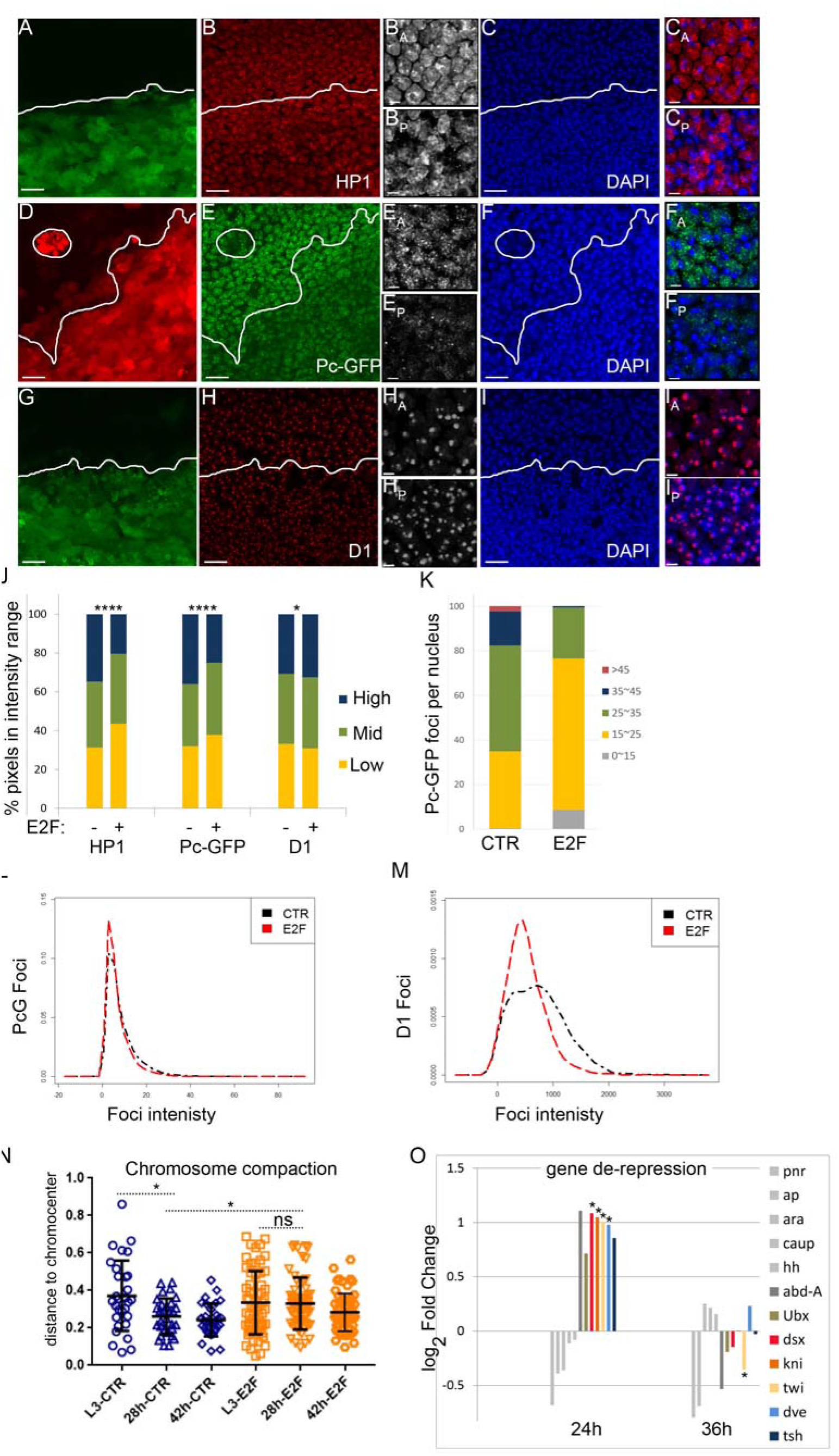
Compromising G0 disrupts D1, HP1 and Polycomb body clustering and leads to partial de-repression of select PcG targets. E2F was co-expressed with GFP or RFP in the posterior wing to delay cell cycle exit. At 26-28h wings were dissected and immunostained for the indicated heterochromatin binding proteins and DAPI to label nuclei (A-I). (J) Overall fluorescence intensities were measured for 319-1270 nuclei for each chromatin modification. The distribution of individual foci intensity (L,M), foci number per nucleus section (K) all indicate that delaying cell cycle exit disrupts heterochromatin clustering and formation of large Polycomb bodies in wing cell nuclei. (N) Chromosome compaction was measured using the distance of the AACAC repeats to the chromocenter. When cell cycle exit is delayed, chromosome compaction is also compromised. For J, N, P-values were determined by an unpaired t-test. *<0.05, **<0.01, ***<0.001, ****<0.0001. (O) Microarray analysis revealed specific PcG target genes that become temporarily de-repressed in wings expressing E2F at 24h when cell cycle exit is delayed. P-values were determined by Anova *<0.05. Scale bars=10 μm in A-I except for anterior (A) and posterior (P) zoomed images where the bar = 2 μm (e.g. BA,BP).

The accumulation of PRC1 components such as Pc into large foci or Pc bodies is important for target gene repression [17, 32]. Since E2F expression disrupts Pc clustering we examined whether increased E2F activity can disrupt the repression of Pc target genes [43–45]. We selected 12 high-confidence Pc target genes predicted not to be direct E2F targets based upon published genome-wide E2F complex binding in *Drosophila* [46]. We examined their expression upon E2F activation in pupal wings at 24 and 36h APF using our previously published array data [47]. We found four Pc targets, *dsx*, *kni*, *twi* and *dve* to be reproducibly de-repressed 1.97-2.12-fold specifically at 24h APF, during the window of time that cell cycle exit is delayed. This suggests that delaying cell cycle exit can partially compromise Pc-dependent gene silencing. E2F activity similarly impacts heterochromatin-dependent gene silencing at the pericentromeric heterochromatin, with the loss of *e2f1* increasing gene silencing by position effect variegation and an increase in E2F activity de-repressing variegated gene expression [48].

In our experiments to delay cell cycle exit E2F is overexpressed for 28h, which includes the final 1-2 normal cell cycles in the pupa wing as well as 1-2 extra cell cycles based upon lineage tracing [18, 47]. We therefore asked whether expression of E2F within only the final cell cycle during terminal differentiation is adequate to disrupt heterochromatin clustering. We used temperature shifts to limit the expression of E2F to a 12h window within the final cell cycle in the pupa and observed a similar disruption of heterochromatin clustering (Supp Fig. 4). We also observed similar effects on heterochromatin clustering when cell cycle exit was delayed by expression of other cell cycle regulators such as CycE/Cdk2 or CycD/Cdk4 (Supp Fig. 5). This demonstrates that heterochromatin clustering in differentiating cells can be disrupted by a single extra cell cycle and that this effect is not specific to E2F overexpression.

### Histone modifications associated with de-condensation are upregulated upon G0 disruption

Compromising G0 leads to the disruption of heterochromatin clustering and chromatin condensation (Fig.4). H3K27ac and H4K16ac are associated with open chromatin such as active enhancers and origins [49–52] and H4K16ac can suppress the formation of higher order chromatin structure [53]. We therefore examined whether these histone modifications were affected by delaying cell cycle exit with E2F overexpression. Indeed during the delay of cell cycle exit, we observed dramatic increase in the levels of these two histone marks throughout the nucleus (Fig. 5A-D). However other histone modifications associated with active chromatin were not affected, such as H3K4me3, pan H3 and H4 acetylation (Fig. 5E-J). Thus, an increase of H3K27ac and H4K16ac could contribute to the compromised chromatin condensation and disruption of heterochromatin clustering observed when cell cycle exit is delayed.

**Fig. 5.**
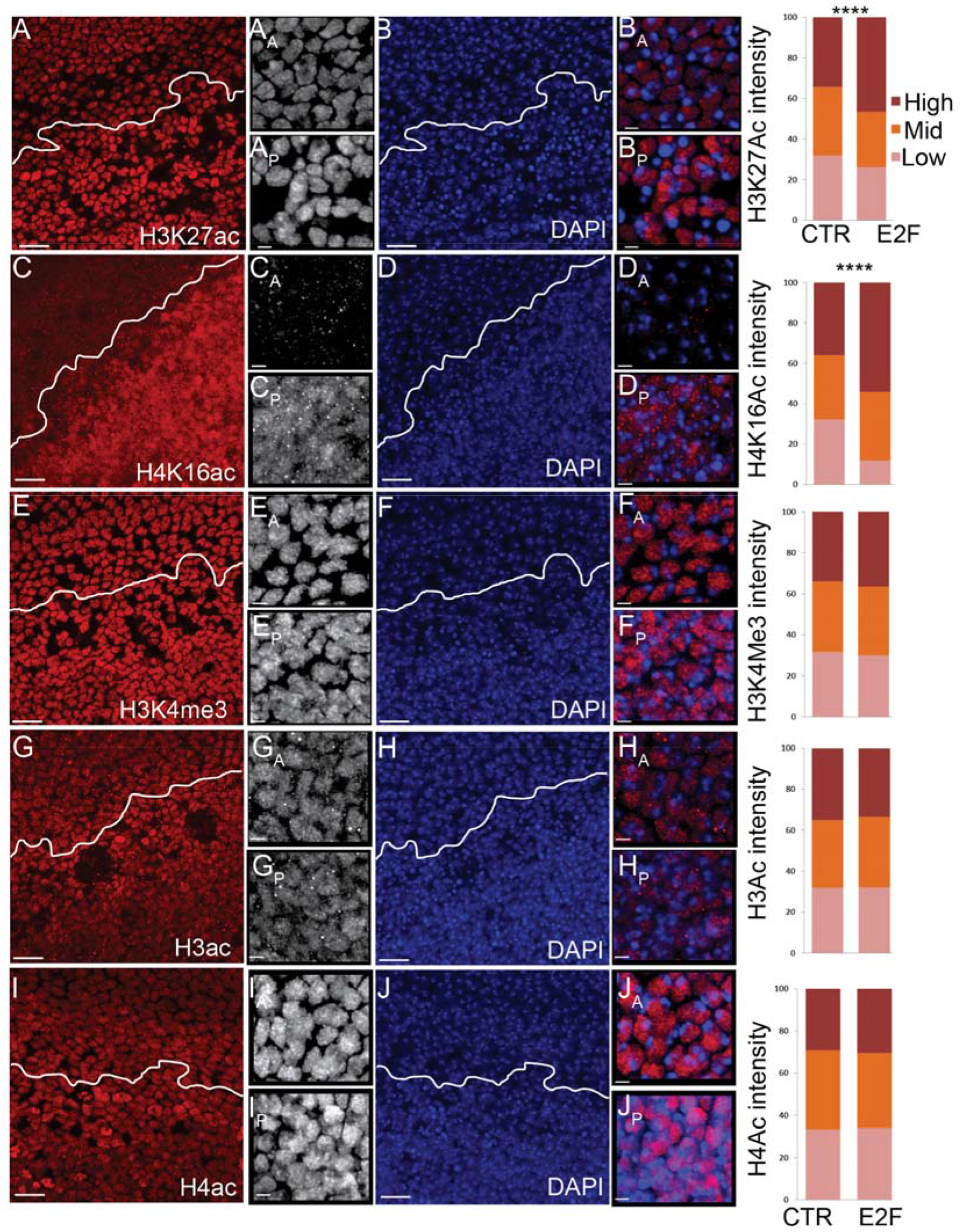
Specific histone modifications associated with gene activation are increased when flexible G0 is compromised. E2F was expressed in the posterior wing to delay cell cycle exit. At 26-28h wings were dissected and immunostained for the indicated histone modifications and DAPI to label nuclei (A-J). The anterior–posterior boundary is indicated by a white line. The distribution of staining intensity in 217-1312 nuclei, binned into three ranges, is shown at right. Compromising flexible G0 specifically increases H3K27ac and H4K16ac. P-values were determined by an unpaired t-test; **** <0.0001. Scale bars=10 μm in A-J except for anterior (A) and posterior (P) zoomed images where the bar = 2 μm (e.g. BA,BP).

### Heterochromatin clustering is restored when cells enter a robust G0 state

Delaying cell cycle exit disrupts heterochromatin clustering, however this is reversible. When we examined wings at 42h -46h, a timepoint when cells enter a robust G0 state refractory to E2F activation, heterochromatin clustering is either partially or completely restored (Fig. 6). Interestingly, levels of H3K27me3 and HP1 became higher in robust G0 after cell cycle exit is delayed (Fig. 6A, 6D). This could be due to the E2F-dependent upregulation of *E(z)* and *Su(var)3-9*, which may indicate an expansion of heterochromatin in differentiating cells that enter a robust G0. Consistent with this idea we also observe an increase in the H2A variant H2Av in E2F-expressing cells in robust G0 (Fig. 6F) and an upregulation of several components of the NuA4 complex responsible for incorporation of H2Av (Supp. Table 1). Heterochromatin expansion is associated with senescence, suggesting delaying cell cycle exit with E2F overexpression could induce oncogenic stress or senescence-like features [54, 55]. Consistent with this, ectopic E2F in the wing induced multiple genes associated with senescence in mammals during robust G0 (Supplemental Table 2) and led to a widespread increase in phosphorylated H2Av, a hallmark of E2F-induced replication stress and DNA damage in *Drosophila* (Fig. 6E) [56].

**Fig. 6.**
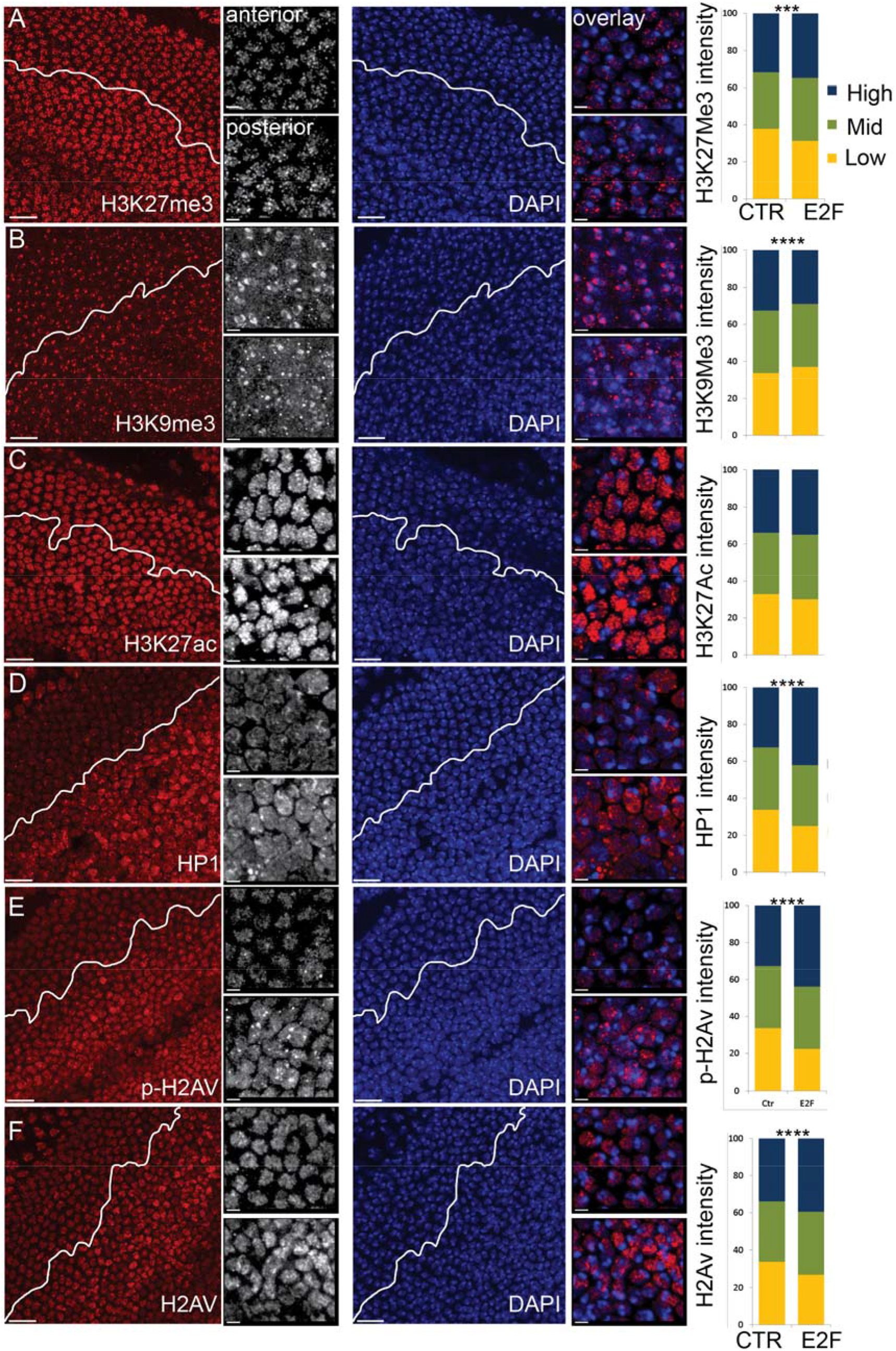
Robust G0 restores heterochromatin clustering and shares features with senescence. E2F was expressed in the posterior wing to delay cell cycle exit. At 42-44h, when cells are in robust G0, wings were dissected and immunostained for the indicated histone modifications or chromatin binding proteins and DAPI to label nuclei (A-F). The distribution of staining intensity in 509-1185 nuclei, binned into three ranges, is shown at right. Robust G0 in the presence of high E2F increases H3K27Me3, HP1 and pH2Av, chromatin marks associated with senescence. P-values were determined by an unpaired t-test; **** <0.0001, ***<0.001. Scale bars=10 μm in A-F except for anterior (A) and posterior (P) zoomed images where the bar = 2 μm (e.g. BA,BP).

### Heterochromatin clustering and cell cycle exit can be uncoupled from terminal differentiation

Heterochromatin clustering becomes restored at the robust G0 phase in the wing as terminal differentiation proceeds. But whether differentiation or cell cycle arrest restores the heterochromatin clustering remains unclear. We previously demonstrated that the robust G0 state in the wing can be bypassed by co-expression of E2F + CycD/Cdk4 [18]. Under these conditions cells in the wing continue cycling past 48h APF, yet physical hallmarks of wing terminal differentiation such as cuticle secretion and wing hair formation proceed after 36h and adult wings form. This condition effectively uncouples cell cycle exit from terminal differentiation in the wing, with actively dividing cells forming actin-rich wing hairs and developing adult cuticle (Fig. 7A-E). We took advantage of this dividing-yet-differentiated context to ask whether heterochromatin clustering requires cell cycle exit. We immunostained 42h wings expressing E2F+CycD/Cdk4 for H3K27Me3, H3K9Me3 and HP1 and found that clustering of facultative and constitutive heterochromatin was dramatically disrupted. We quantified facultative heterochromatin foci and found H3K27Me3 forming fewer, smaller and less intense foci (Fig. 7 F-K). By contrast, HP1 levels became extremely high, with a diffuse localization throughout the nucleus (Fig. 7H,J), similar to the effects of E2F on HP1 at robust G0 (Fig. 6). These results demonstrate that heterochromatin clustering is a consequence of cell cycle exit rather than terminal differentiation. In addition, terminal differentiation can proceed despite a visibly significant disruption of heterochromatin organization.

## Discussion

### The relationship between heterochromatin clustering and differentiation

A number of studies have documented increased clustering and condensation of heterochromatin as cells differentiate [reviewed in 57]. In this study we reveal a substantial effect of the cell cycling status on heterochromatin clustering independent of differentiation. Heterochromatin clustering increases as the cell cycle slows and cells exit the cell cycle. By delaying or bypassing cell cycle exit in terminally differentiating cells, we show that the highly clustered state of heterochromatin in postmitotic cells is a consequence of cell cycle exit rather than the process of terminal differentiation.

Importantly, we show that differentiation still proceeds even when cell cycle exit is prevented and heterochromatin clustering is severely disrupted (Fig.7). We suggest this is because disrupting heterochromatin clustering has only limited effects on the expression of specific heterochromatin-repressed genes in the context of the *Drosophila* wing (Fig. 4) and minimal effects on the terminal differentiation gene expression program. Indeed, we show that cell cycle exit can proceed normally in the *Drosophila* wing even when heterochromatin-dependent gene silencing is directly compromised (Fig. 2). Altogether this demonstrates that the increased heterochromatin clustering observed during differentiation is a consequence rather than a cause of cell cycle exit and raises questions regarding the function of increasing very long-range heterochromatin interactions and heterochromatin clustering in differentiation.

### What is the function of heterochromatin clustering?

When we delay or bypass cell cycle exit, we visibly disrupt heterochromatin clustering, however local heterochromatin clustering likely remains intact. We find that this leads to very mild effects on the expression of only a small number of Polycomb target genes (Fig.4), and we did not find significant de-repression of genes that are located in or near constitutive heterochromatin [58] (not shown). Our result is consistent with recent work showing that compromising some types of PcG clustering seems to have limited and selective effects on Polycomb target gene silencing [32]. However, the minimal effect of disrupting cell cycle exit on heterochromatin-dependent gene silencing is somewhat unexpected as the E2F1 gene was one of the early-identified modifiers of position-effect variegation (PEV), which is thought to be due to heterochromatin-dependent gene silencing through association with constitutive heterochromatin [48, 59, 60]. This suggests either the PEV assay is highly sensitive to even mild or selective changes in heterochromatin-dependent gene silencing or that this assay reads out changes in the chromatin state that are different from the silencing of the endogenous genes we examined. Indeed, there are additional possible functions for heterochromatin clustering beyond heterochromatin-dependent gene silencing. For example, heterochromatin clustering could facilitate DNA damage repair in postmitotic cells, which downregulate many DNA repair genes when they exit the cell cycle and become more reliant on error prone NHEJ [reviewed in 61]. Sequestration of heterochromatin may prevent inappropriate interactions and fusions. It has also been proposed that sequestration of heterochromatin could lead to an increased efficiency of gene activation for very highly expressed genes by reducing the availability of possible binding sites for specific transcription factors [57], or could facilitate the formation of transcription factories [62].

A number of other studies also describe changes in the abundance of specific chromatin modifications associated with entry into or exit from G0. For example, H3K9Me3 and H3K27Me3 accumulate in postmitotic, differentiated cardiac muscle [38] while H4K16Ac and H3K27Ac increase in activated B cells exiting G0 [63]. While our data from the *Drosophila* wing suggests clustering of H3K9Me3 and H3K27Me3 domains during cell cycle exit rather than obvious changes in total levels, we do observe a strong up-regulation of H4K16 acetylation when G0 is delayed by E2F activation, a situation similar to cell cycle re-entry from G0. We also observe a decrease in H4K20Me3 when G0 is compromised, similar to what has been reported for quiescent human fibroblasts [64]. While H4K16Ac was not specifically measured in the fibroblast study, H4K20 methylation and H4K16 acetylation are antagonistic marks [65] and H4K16 acetylation can decompact nucleosomes *in vitro*, although whether this also occurs *in vivo* has been questioned [53, 66]. We suggest some aspects of chromatin remodeling, such as compaction and coalescence of heterochromatin (which may be tied to H4K16/20 dynamics) are shared among different contexts of cell cycle exit/re-entry, while other chromatin changes associated with G0 entry/exit may be more cell type specific.

### Why does delaying or bypassing cell cycle exit disrupt heterochromatin clustering in interphase?

Our experiments effectively separate cell cycle exit from terminal differentiation to reveal that heterochromatin clustering is a consequence of cell cycle exit. Furthermore, heterochromatin clustering can be disrupted within a single cell cycle (Supp. Fig 4), suggesting progression through one round of S or M-phase is sufficient to disrupt heterochromatin clustering and long-range interactions. The effects we observe are not due to the dilution of chromatin marks by incorporation of new histones in S-phase, since we do not see changes in all histone marks (e.g. H3K4 methylation) or reduced global levels of histone marks in proliferating vs. postmitotic cells (Supp. Fig. 1). Indeed, when global levels of chromatin marks in actively proliferating fibroblasts were quantified and compared to fibroblasts held in G0 under contact inhibition for 14d, the majority of histone modifications did not exhibit significantly different levels [64]. This is likely because the levels of many histone modifiers are upregulated by positive cell cycle regulators through E2F transcriptional activity (Supp. Table 1) which effectively coordinates increased histone modification with increased production and incorporation in S-phase.

Overexpression of E2F could have effects on chromatin modifications and condensation through sequestration or indirect inhibition of RB-family proteins via increased Cyclin/Cdk expression. RB associates with chromatin modifying complexes that promote facultative and constitutive heterochromatin formation [67–69]. RB also impacts chromosome condensation and cohesin levels at pericentromeric heterochromatin [70–72]. However in our experiments during robust exit, E2F levels and transcriptional targets remain high while heterochromatin clustering and chromatin marks are restored (Fig.6). This suggests that even if RB is inhibited by overexpression of E2F, the eventual entry into robust G0 somehow restores heterochromatin organization and chromatin modifications independent of RB.

During mitosis most transcription factors and chromatin modifiers are ejected from chromatin and higher order architecture is lost [7]. This together with mitotic spindle assembly leads to the loss of long-range interactions and interchromosomal associations. These interactions are then restored, even in the presence of high E2F activity once cells engage additional mechanisms to exit the cell cycle during the robust G0 phase [47]. Our findings are in agreement with previous studies showing that the motion of heterochromatin domains and Polycomb bodies become more constrained as the cell cycle slows and cells exit the cell cycle [16, 17]. We suggest that constrained motion combined with increased self-association or polymerization likely leads to the coalescence of heterochromatin after cell cycle exit.

## Conclusions

Heterochromatin clusters as cell exit the cell cycle and terminally differentiate. Delaying or preventing cell cycle exit disrupts heterochromatin clustering and globally alters chromatin modifications. Heterochromatin clustering during terminal differentiation is a consequence of cell cycle exit, rather than differentiation. Compromising heterochromatin-dependent gene silencing does not disrupt cell cycle exit.

## Acknowledgements

A special thank you to Abbey Roelofs (University of Michigan, Advocacy and Research Support, LSA-IT) for developing the automatic quantification toolkit in Matlab. We thank Dr. Yukiko Yamashita (University of Michigan, Ann Arbor) for sharing FISH reagents and protocols as well as D1 antibody. We thank Dr. Keith Maggert (Texas A&M University) for sharing the Y10C reporter and Dr. Terry Orr-Weaver (Massachusetts Institute of Technology) for sharing the E2F1 antibody. Additional antibodies were obtained from Developmental Studies Hybridoma Bank (DSHB), created by the NICHD of the NIH and maintained at The University of Iowa. Stocks obtained from the Bloomington Drosophila Stock Center (NIH P40OD018537) were used in this study. The microarray data referenced has been deposited in GEO, accession number GSE30484. This work in the Buttitta Lab was supported by the NIH (R21AG047931) and the American Cancer Society (RSG-15-161-01-DDC). Y.M performed all experiments. Y.M. and L.B. conceived of the project, analyzed the data and wrote the manuscript. The authors have no conflicts of interest.

---

**Fig. S1.**
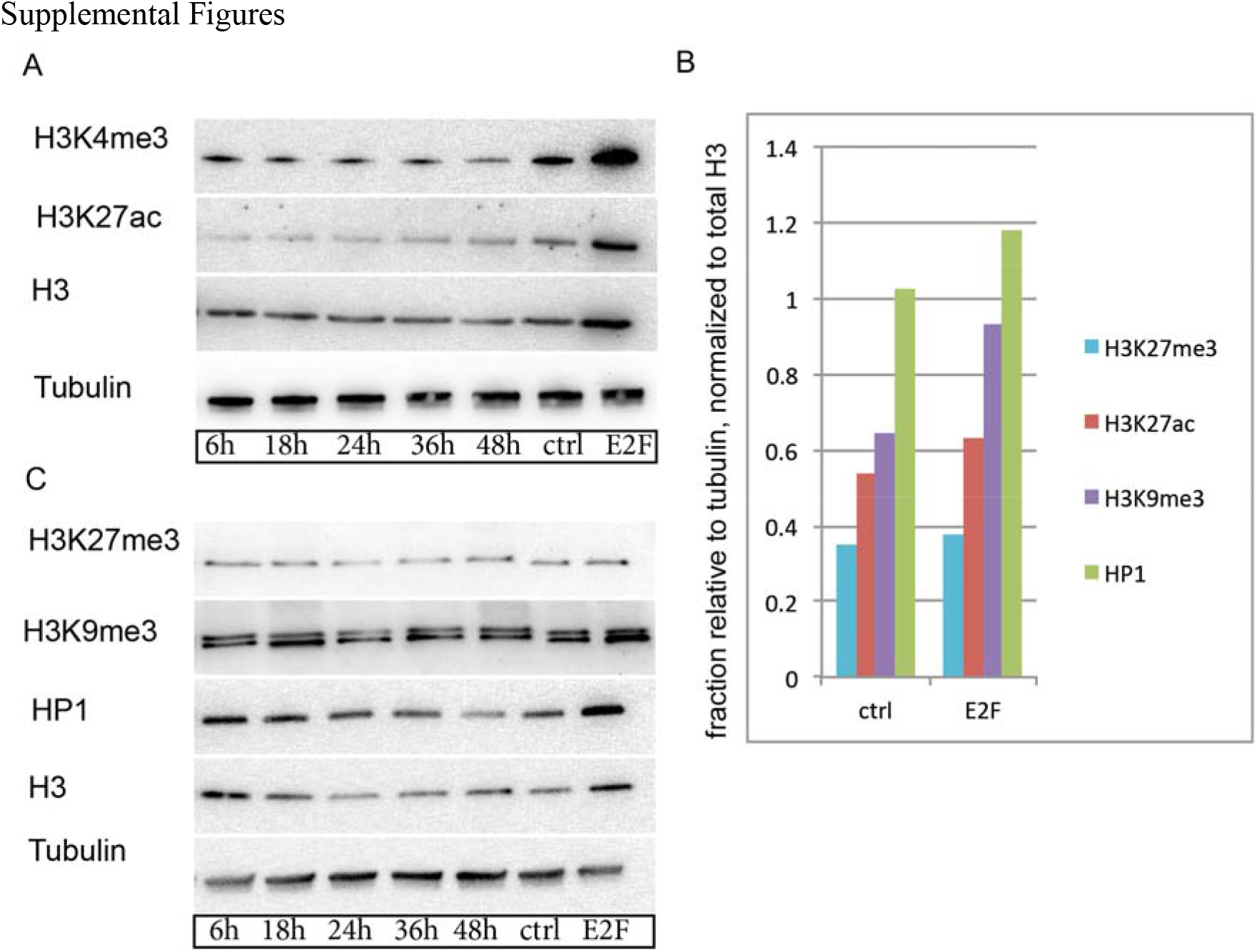
Global levels of histone modifications do not dramatically change at cell cycle exit. (A-D) Quantitative western blots were performed on wings of the indicated stages to assess the levels of modified or total histone H3 or HP1. Control (Ctrl) and E2F samples are from 28h postmitotic wings overexpressing GFP or E2F respectively. Total H3K9Me3, H3K27Me3, and HP1 levels do not dramatically change with cell cycle exit, however they increase with E2F expression. Modifications associated with active chromatin, H3K4Me3 and H3K27Ac also do not dramatically change with cell cycle exit, but increase upon E2F expression.

**Fig. S2.**
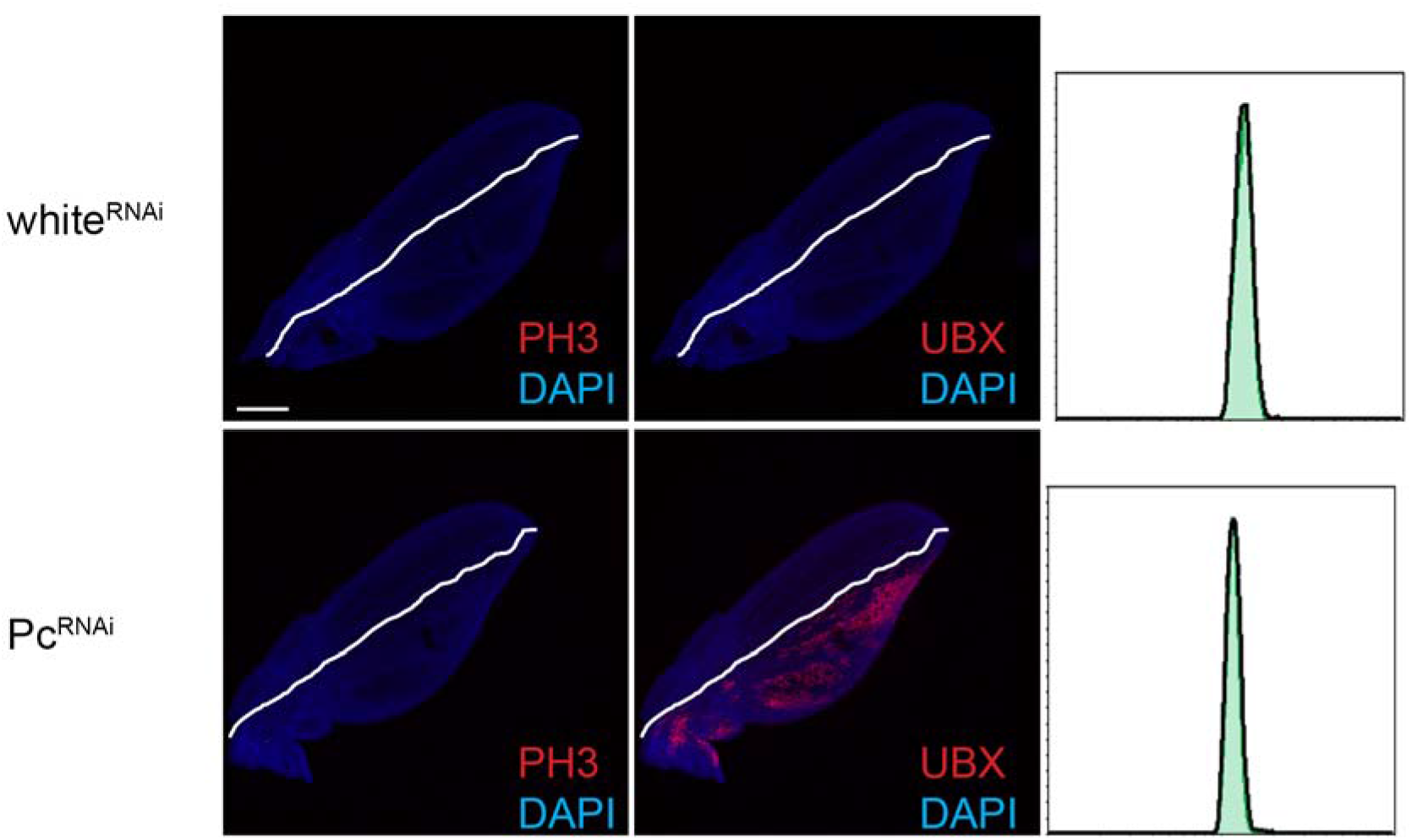
Compromising PRC1 does not delay cell cycle exit. RNAi to *Pc* or *white* (as a control) was expressed in the posterior wing from the L3 stage and postmitotic wings at 26-28h were examined for mitoses as indicated by PH3 and effective knockdown of PRC1 function by de-repression of the PRC1 target gene Ubx. Flow cytometry was also performed to measure cells that enter S and G2 phases. Green trace indicates cells from the posterior wing expressing the indicated transgenes. Black trace: control non-expressing anterior wing cells. Compromising PRC1 activity does not delay cell cycle exit. Scale bars=100 μm.

**Fig. S3.**
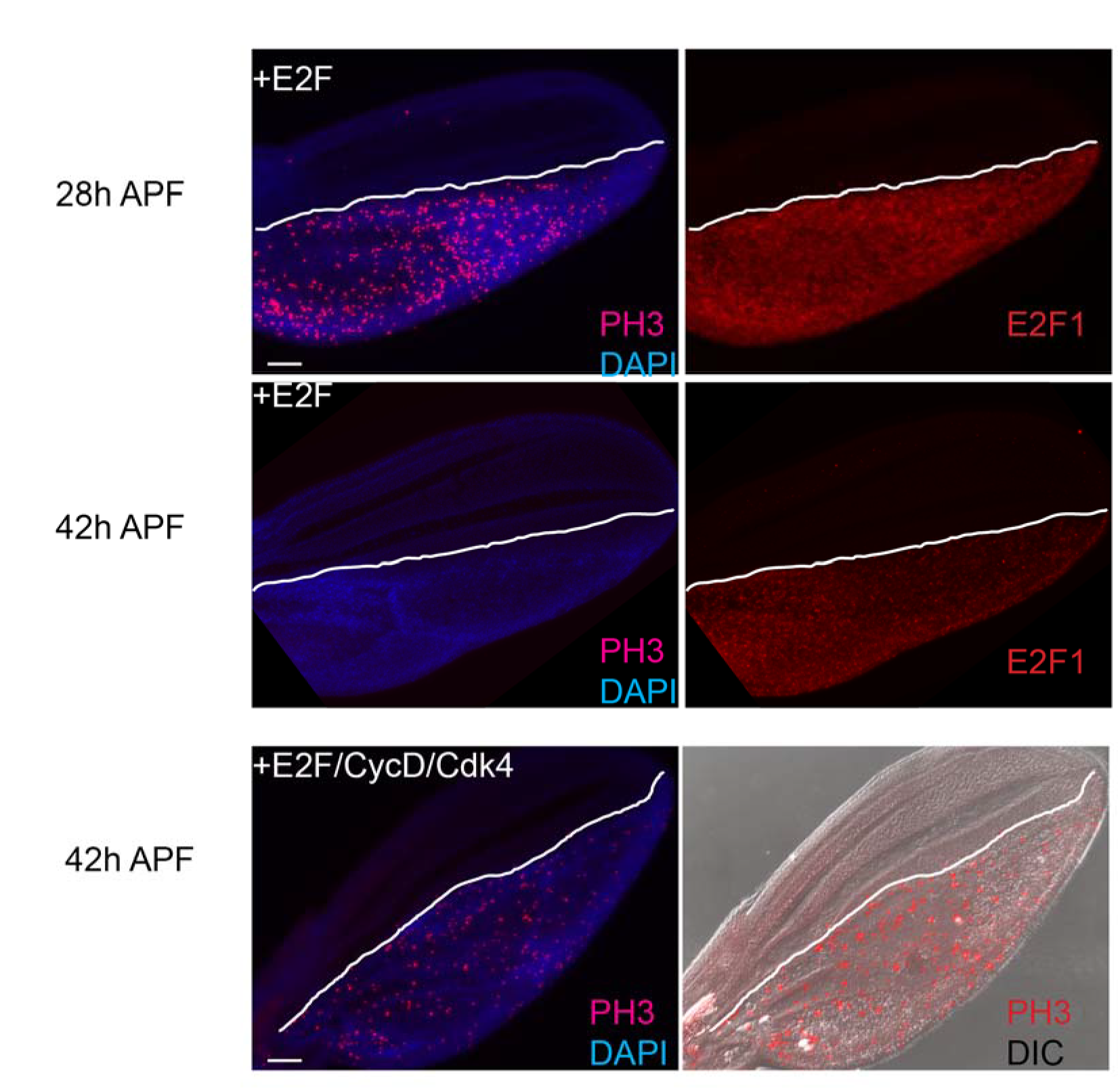
Two stages of G0 in differentiating wings. E2F was expressed in the posterior wing to delay cell cycle exit. 28h and 42h APF pupal tissues were dissected and immunostained for PH3 (to label mitoses) and E2F1. The anterior/posterior boundary is specified by the white line. Overexpression of E2F delays entry into G0 until 36h. At 42h cells expressing high E2F1 are postmitotic (in robust G0). CycD/Cdk4+E2F expression in the posterior wing is able to bypass the robust G0 to promote continued cycling, as shown by abundant mitoses (PH3) at 42h. Bar= 50μm

**Fig. S4.**
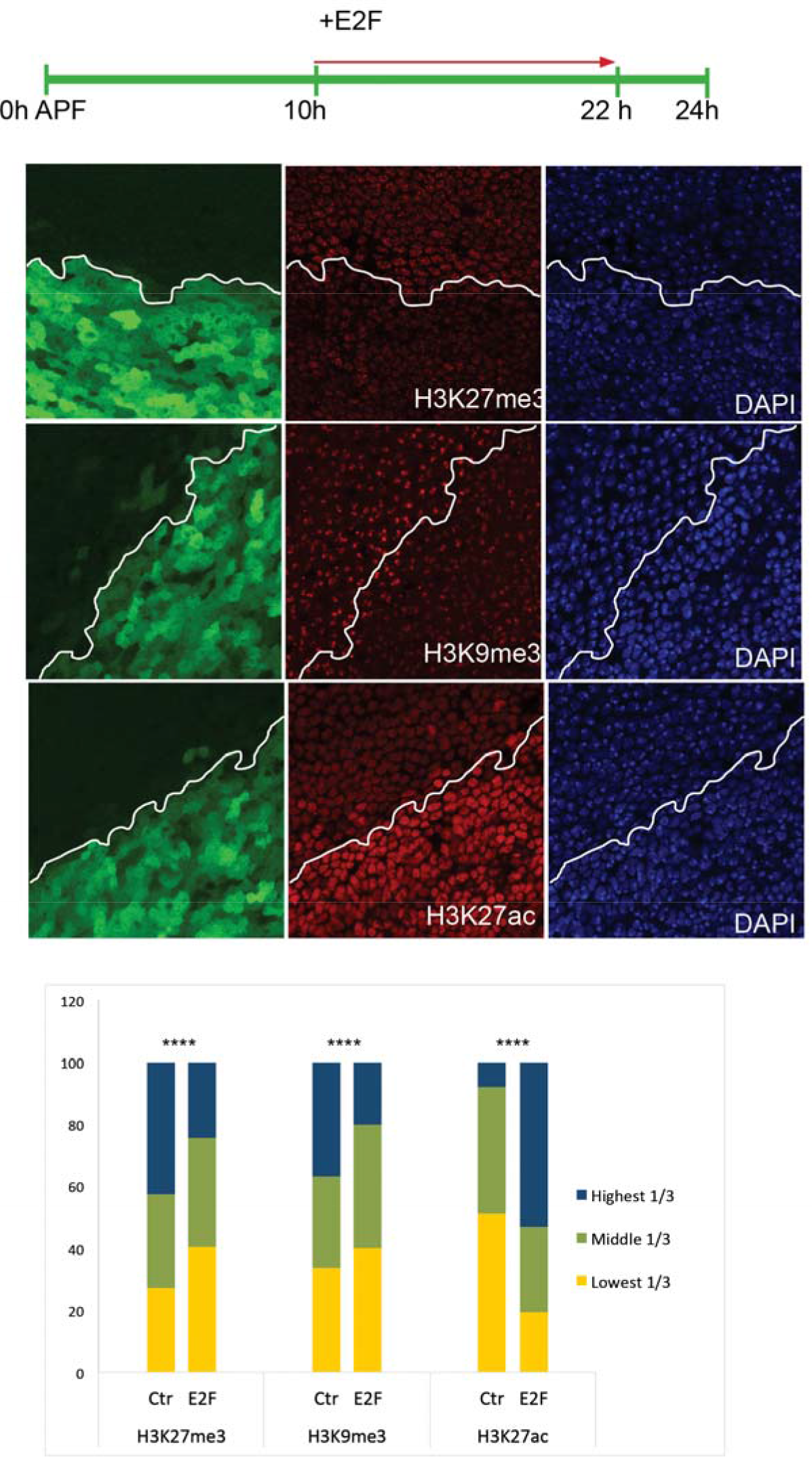
Clustering of heterochromatin can be disrupted within one cell cycle. E2F was overexpressed in the posterior wing from 10h APF. 12h later (within approximately one cell cycle) tissues were immunostained for indicated histone modifications. The posterior region is labeled by the expression of GFP and the anterior/posterior boundary is specified by the white line. The distribution of staining intensity in 1112-1339 nuclei, binned into three ranges, is shown at bottom. E2F disrupts heterochromatin clustering within one cell cycle. P-values were determined by an unpaired t-test; **** <0.0001.

**Fig. S5.**
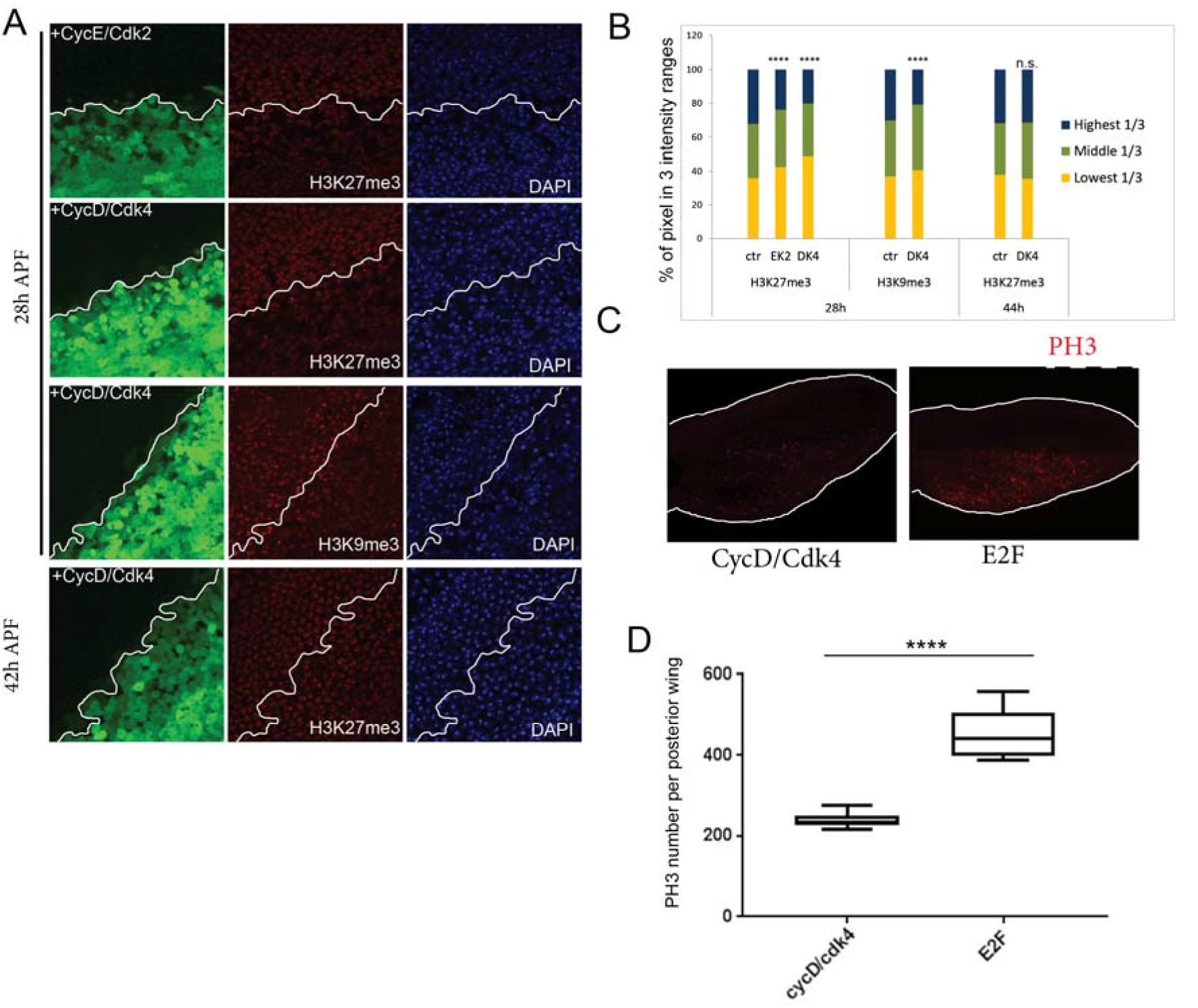
Delaying cell cycle exit disrupts heterochromatin. (A) CycE/Cdk2 or CycD/Cdk4 complexes were overexpressed in the posterior wing from 0h APF. The anterior/posterior boundary is indicated by the white line. At 28h (flexible G0) or 42h APF (robust G0) pupal tissues were dissected and immunostained for the indicated histone modifications. (B) The distribution of staining intensity from 492-976 nuclei, binned into three ranges, is shown. Wings expressing E2F or CycD/Cdk4 to delay cell cycle exit were stained for mitoses (PH3) and the mitotic index at 27h was quantified for the posterior compartment (C-D). The degree of heterochromatin disruption correlates with the number of cells cycling. P-values were determined by an unpaired t-test; ****P-value <0.0001.

**Supplemental Table 1.**
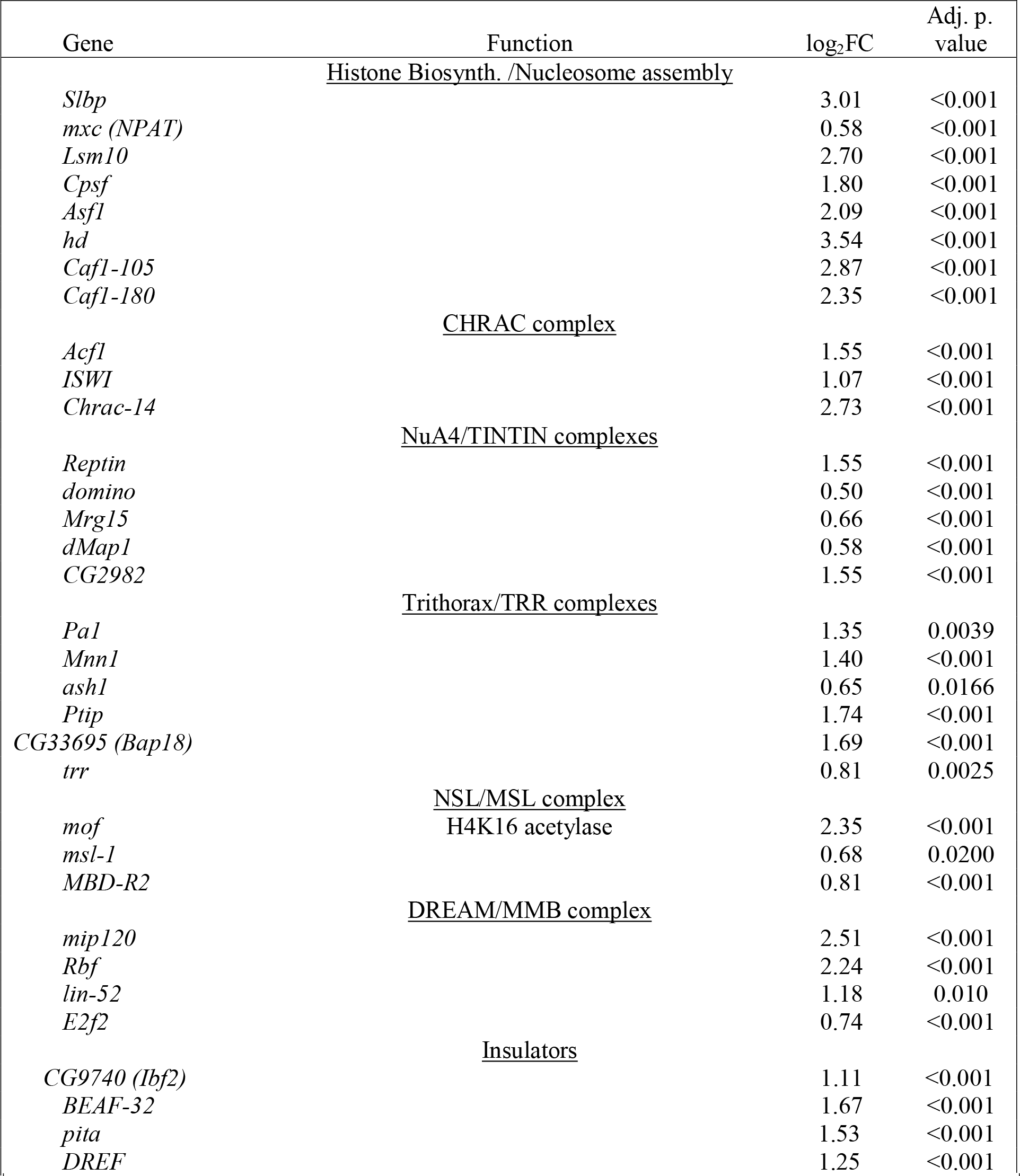
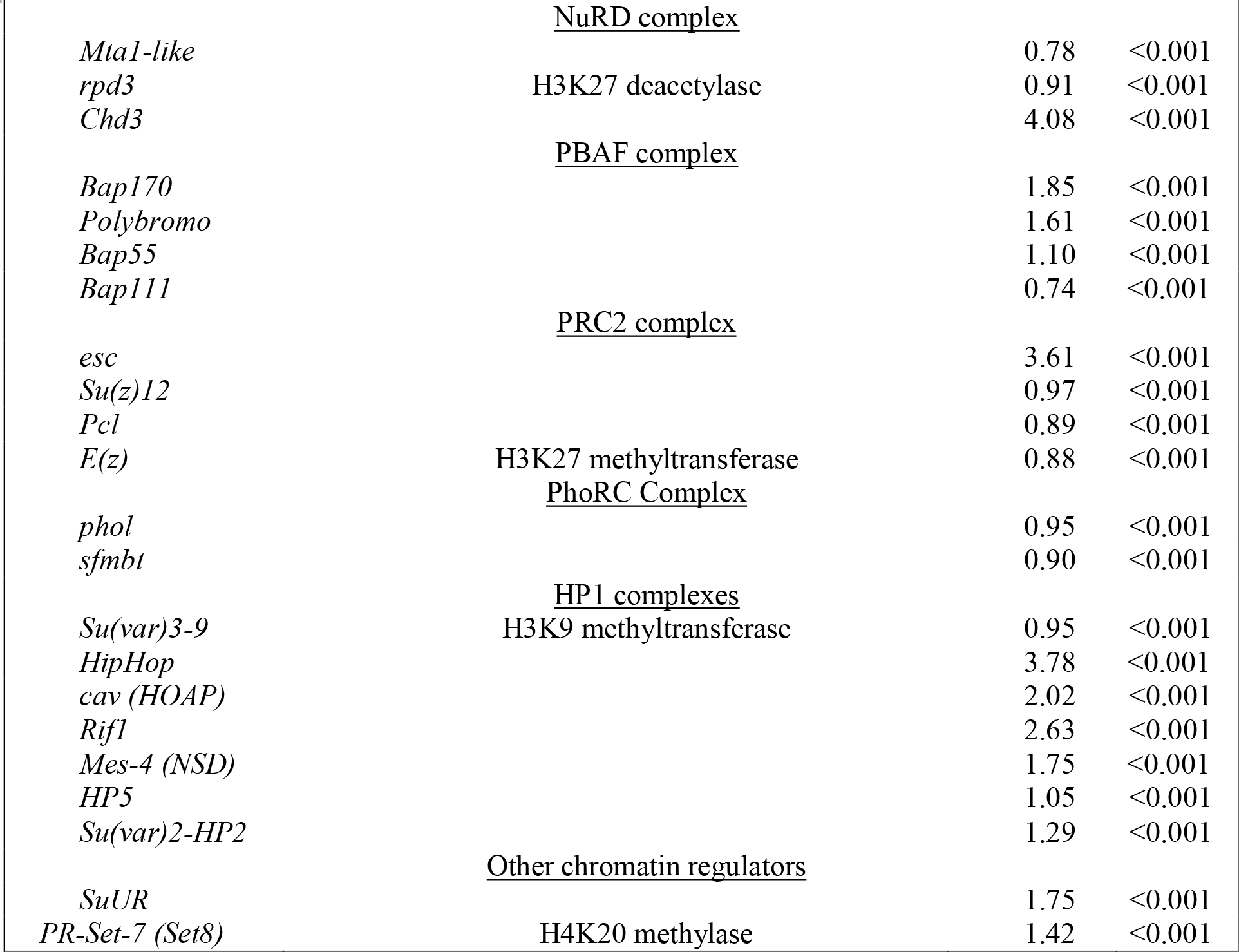
Chromatin modifiers/organizers/remodelers that are upregulated upon E2F1/DP expression in pupal wings

**Supplemental Table 2.**
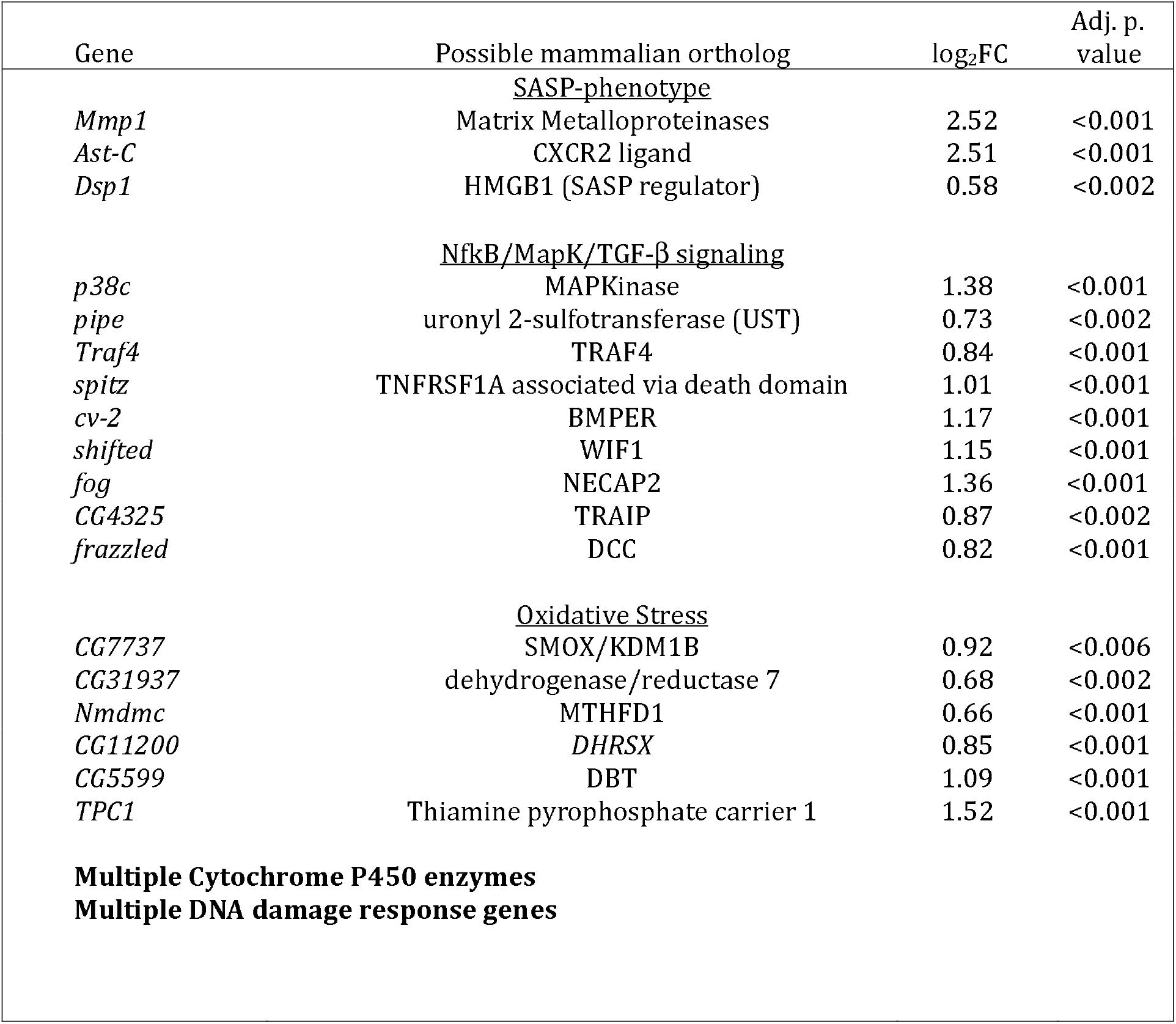
Genes associated with senescence that are upregulated during robust G0 in the presence of ectopic E2F1/DP.

